# Mid-level visual features engage V4 to support sparse object recognition under uncertainty

**DOI:** 10.1101/2025.10.28.685033

**Authors:** Leyla Loued-Khenissi, Elsa Scialom, Antoine Lutti, Michael Herzog, Bogdan Draganski

## Abstract

Cortical visual prostheses aim to restore sight by stimulating the visual cortex, yet current approaches remain limited. Implants targeting V1 typically evoke isolated phosphenes that do not support coherent object recognition. We asked whether mid-level visual features, specifically curve segments associated with area V4, could provide a more efficient substrate for artificial vision. Using fMRI, 24 participants performed an object recognition task with stimuli fragmented into either phosphene-like dots or curve segments at varying densities. Behaviorally, curve segments enabled recognition at lower density thresholds and produced steeper psychometric slopes. Neurally, both fragment types engaged V1, but only curve segments recruited V4, and correct recognition of segments relative to phosphenes evoked broader occipito-temporo-parietal responses. Multivariate decoding showed that successful recognition of phosphenes was predicted by early visual and cingulate activity, whereas recognition of segments was predicted only by V4, indicating a shift toward mid-level representational support. Mediation analyses further showed that phosphene-based recognition relied on anterior cingulate cortex under conditions of higher uncertainty, while segment-based recognition depended primarily on visual cortex. Together, these results show that mid-level fragments confer perceptual and neural advantages, reducing reliance on both dense stimulation and downstream uncertainty resolution, and identify V4 as a promising target for next-generation cortical prostheses.

## Introduction

Vision is one of the most important human faculties, yet approximately 40 million people worldwide suffer from severe visual impairments that significantly reduce quality of life. Retinal prostheses have shown partial success in specific degenerative conditions, but their performance is constrained by the dependence on residual retinal circuitry and coarse resolution (Erickson-Davis & Korzybska, 2021). This has motivated a long-standing interest in cortical visual prostheses, which bypass the eye to stimulate visual cortex directly. Electrical stimulation of the occipital lobe has been known for over a century to induce phosphenes — points of light perceived in the absence of external input — confirming the basic plausibility of such an approach (S. C. Chen et al., 2009).

Despite this conceptual promise and ongoing clinical trials (Fernández et al., 2021), no safe and reliable cortical prosthesis has been realized to date. Current devices face numerous technical and biological challenges (Borda & Ghezzi, 2022). Compared with cochlear implants, which act on an accessible single nerve, cortical devices require invasive neurosurgery and must operate with relatively few stable electrodes (Liu et al., 2022). In addition to technical limitations, the gap between perceptual promise and functional vision reflects a deeper representational problem: current implants allow too few stimulations too early, based on neural codes that are too simple. Targeting V1 has been the default strategy because stimulation at this site reliably produces phosphenes. However, because each electrode produces a unique point of light, reconstructing coherent shapes requires a prohibitively high electrode density. In macaque studies, small-scale intracortical stimulation in V1 evoked stable, spatially specific percepts over years (Davis et al., 2012), whereas higher-density Utah array implants showed a reliable initial performance but rapid degradation of signal quality and perceptual stability over time (Chen et al., 2020), highlighting a trade-off between channel count and long-term functional reliability. In humans, multi-electrode stimulation of early visual areas produces spatially organized but unstable percepts: simultaneous stimulation in V1–V3 evokes multiple phosphenes that rarely merge into coherent forms (Bosking et al., 2022) and perceptual quality varies widely with training, fixation, and electrode mapping (Oswalt et al., 2021). Together, these results indicate that while direct cortical stimulation can elicit basic visual sensations, a “connect-the-dots” approach yields unstable, fragmentary percepts rather than meaningful vision (Sanchez-Garcia et al., 2018).

One avenue to explore in addressing the sparse phosphene problem is to determine how mid-level visual organization can be leveraged to encourage a coherent percept from fragmented input. One possible solution is to move beyond V1 and target downstream areas that encode mid-level visual features. Rather than targeting the building blocks of vision in V1, an alternative is to engage the brain’s intrinsic feature-building machinery, the mid-level codes that bind local edges into global form. Area V4 is an especially promising candidate for this search as it is tuned to curved segments and intermediate shape features (Carlson et al., 2011; Wei et al., 2018), and contributes directly to object recognition (Roe et al., 2012). Human neuroimaging and retinotopic mapping confirm that V4 contains curvature-selective domains and occupies a pivotal position along the ventral visual pathway (Winawer & Witthoft, 2015). Psychophysical and fMRI evidence further shows that contour integration and curvature processing emerge through V4 and strengthen toward lateral occipital cortex, linking mid-level structure to global shape perception (Kuai et al., 2017; Lawrence et al., 2023). These findings align with classic recognition-by-components theory that emphasizes curvature and collinearity as litmus tests for object recognition (Biederman, 1987). Representations at this “mid-level” stage in the ventral visual stream may provide a more efficient basis for percept formation than discrete point-like inputs.

There have been very limited studies investigating V4 stimulation’s effect on vision. Shiozaki and colleagues (Shiozaki et al., 2012) showed that microstimulation of disparity-selective clusters in macaque V4 biases depth judgments, establishing a causal role for V4 neurons in depth discrimination. Another study in macaques found V4 microstimulation had virtually no effect on visual stimulus detection thresholds (Dagnino et al., 2015) but a more recent study found microstimulation of area V4 in macaques led to improved performance in a visual detection task (Kienitz et al., 2022). Yet it remains unknown how V1 and V4 contribute to recognizing objects from sparse, fragmented representations composed of either low-level points or mid-level segments. In other words, does the brain’s reliance on mid-level codes for natural vision also determine its success under sparse perceptual information, the regime most relevant for prosthetic sight? Although this possibility has been raised in animal studies, it remains largely unexplored in humans. Moreover, no study has directly tested whether the perceptual benefits of contour continuity, a feature central to mid-level visual organization, can be traced to V4 processing, which may also ultimately support more coherent cortical prosthetic vision.

By integrating evidence from cortical prosthesis research and mid-level vision, we examine whether the perceptual gains afforded by contour continuity originate from mid-level curvature and grouping mechanisms in area V4, identifying a potential, alternative stinulation site for more coherent prosthetic implantation. Here, we addressed this question using fMRI and a task designed to simulate recognition from prosthesis-like input. Participants viewed fragmented outlines of objects composed of either phosphene-like dots or curve segments at increasing density levels, and then later identified the object from a choice grid. This design allowed us to test three specific hypotheses. First, we hypothesized that curve segments would facilitate more efficient recognition than phosphenes, requiring fewer fragments for successful object identification (H1). Second, we hypothesized that successful recognition from curve segments would preferentially engage mid-level extrastriate cortex, specifically V4, in contrast to phosphenes, which would rely more heavily on V1 (H2). Finally, we hypothesized that correct recognition under sparse and uncertain conditions would recruit the anterior cingulate cortex (ACC), a downstream control region involved in uncertainty resolution (Levinson et al., 2021). We further predicted that this contribution would be stronger for phosphenes, which provide less reliable sensory information, than for curve segments (H3). By identifying how the brain reconstructs objects from sparse cues, this study bridges visual neuroscience and neuroengineering, offering representational principles to guide the next generation of cortical visual prostheses.

## Results

### Behavioral

To examine specific differences between fragments types, our main area of interest, the following results reflect only fragmented image trials.

To determine if object recognition occurs at a lower density threshold with segments rather than phosphenes, we computed individual subjects proportion of correct responses for each density level for each fragment type, and fit a logistic model to these values of the form

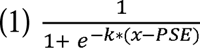

This fit provides the *k* parameter, the steepest slope of the curve, indicative of a transition between incorrect and correct; and the points of subjective equality (PSE), indicating the density level at which this transition occurs. We find, for segments, a mean of PSE = 31.74, (SD = 18.60) and for phosphenes, a mean PSE of = 43.44 (SD = 13.09). A direct comparison of these fits show a significant difference (segments < phosphenes): t =-5.35, p < 0.001 (Figure 1a).

The mixed-effects regression model assessing factors influencing accuracy shows significant positive effects of segment fragment type (t = 4.110, p < 0.001) and density (t = 13.314, p < 0.001). Accuracy is negatively associated with reaction time (t = -4.444, p < 0.001). In addition, accuracy is reduced by the interactions of segment fragment type with density (t = - 3.479, p < 0.001) and with entropy (t = -2.591, p = 0.010). The effect of entropy alone is nonsignificant (t = 0.140, p = 0.888), as is the interaction of segment fragment type with reaction time (t = -0.753, p = 0.451) (Figure 1b).

The mixed-effects regression model assessing factors influencing confidence ratings shows significant positive effects of accuracy (t = 11.809, p < 0.001) and density (t = 12.901, p < 0.001). Confidence is negatively associated with entropy (t = -5.760, p < 0.001). The main effect of segment fragment type is nonsignificant (t = -0.230, p = 0.818), as are the effects of confidence reaction time (t = -0.943, p = 0.346) and the interactions of segment fragment type with density (t = 0.910, p = 0.363), accuracy (t = 0.245, p = 0.806), confidence reaction time (t = -0.921, p = 0.357), and entropy (t = 1.110, p = 0.267) (Figure 1).

**Figure 1:**
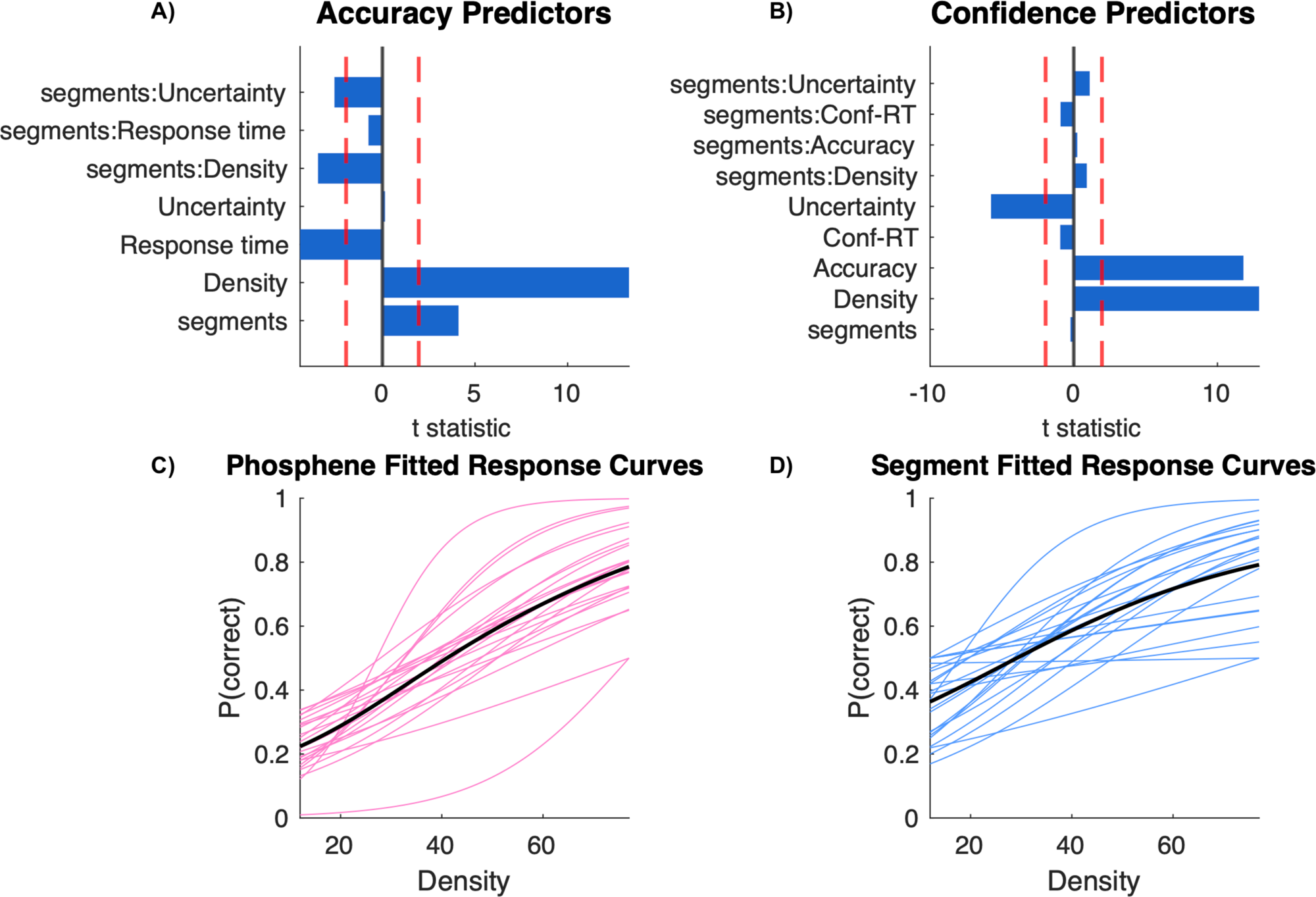
Behavioral analysis of responses. a) and b) T statistics for different predictors of accuracy and confidence. Red dashed lines indicate significance thresholds for p = 0.05. c) and d) Psychometric curves for phosphene and segment trials fit to a logistic function. Pink curves indicate individual subject performance in phosphene trials against fragment density levels. Blue curves indicate individual subject performance in segment trials against fragment density levels. The black curve indicates the grand mean fit.

Taken together, the hypothesis that segment fragment type and density enhance object recognition were confirmed. These hypotheses are further edified by the positive effects of accuracy and density on metacognitive confidence, which provide evidence that individuals are aware of their performance quality.

### Neuroimaging - Whole brain results

#### Control Trials: Contour and RGB Trial Tests at Stimulus Presentation

RGB statistical maps showed no significant responses. Contour image trials yielded clusters in the bilateral inferior temporal gyri, fusiform gyri, left angular gyrus, left hippocampus and bilateral inferior frontal gyrus. Contour trials tested as positive against all fragment types showed clusters bilaterally in the fusiform gyrus (as did contour trials tested against segment trials) and clusters in the left fusiform and right inferior temporal gyrus in phosphene trials. All fragmented image trials tested as greater than contour trials yielded a widespread network of clusters in the following regions: bilateral precentral gyrus, left supplementary motor cortex, right cerebellum, left parietal operculum, left angular gyrus, left thalamus, left putamen, left cuneus, bilateral middle frontal gyrus, left inferior occipital gyrus, left cuneus, left mid cingular gyrus, right lingual gyrus, and right postcentral and supramarginal gyri.

Phosphene trials greater than contour trials showed clusters in right precentral gyrus and right inferior occipital gyrus. Segment trials greater than contour trials showed a more extensive pattern of clusters in left supramarginal and superior temporal gyri, as well as right lingual gyrus. With respect to contour trials greater than RGB, numerous clusters emerged including: bilateral inferior, middle temporal and fusiform gyri, right parahippocampal and angular gyri, and left anterior insula. No other tests on RGB trials yielded significant clusters.

#### Phosphene and Segment Trial Tests

Phosphene trials at cue presentation onsets yielded bilateral clusters in the middle temporal gyrus, left angular gyrus and extensively in the cingulate (Figure 2a and 2b). Segment trials yielded significant clusters in similar regions, as well as in the thalamus. Phosphene density was associated with significant bilateral responses in the fusiform gyrus, while segment density was associated with right mid and left inferior occipital cortex (Figure 2c and 2d). Phosphene confidence was associated with left inferior occipital cortex, while segment confidence showed a similar pattern of response that did not survive correction for multiple comparisons (Figure 2f)(Table 2). When testing phosphene versus segment contrasts directly, only one test yielded significant results, at response onsets: correct segment trials greater than correct phosphene trials showed significant results in the cerebellum, middle temporal lobe, bilateral supramarginal and postcentral gyri, middle and anterior cingulate, as well as the left temporal pole and left anterior insula, among other regions (Table 3). Critically, the cluster with a peak in the cerebellum extends over area V4 on the right (Figure 3b). No other tests examining phosphene versus segment trials yielded significant results.

**Figure 2.**
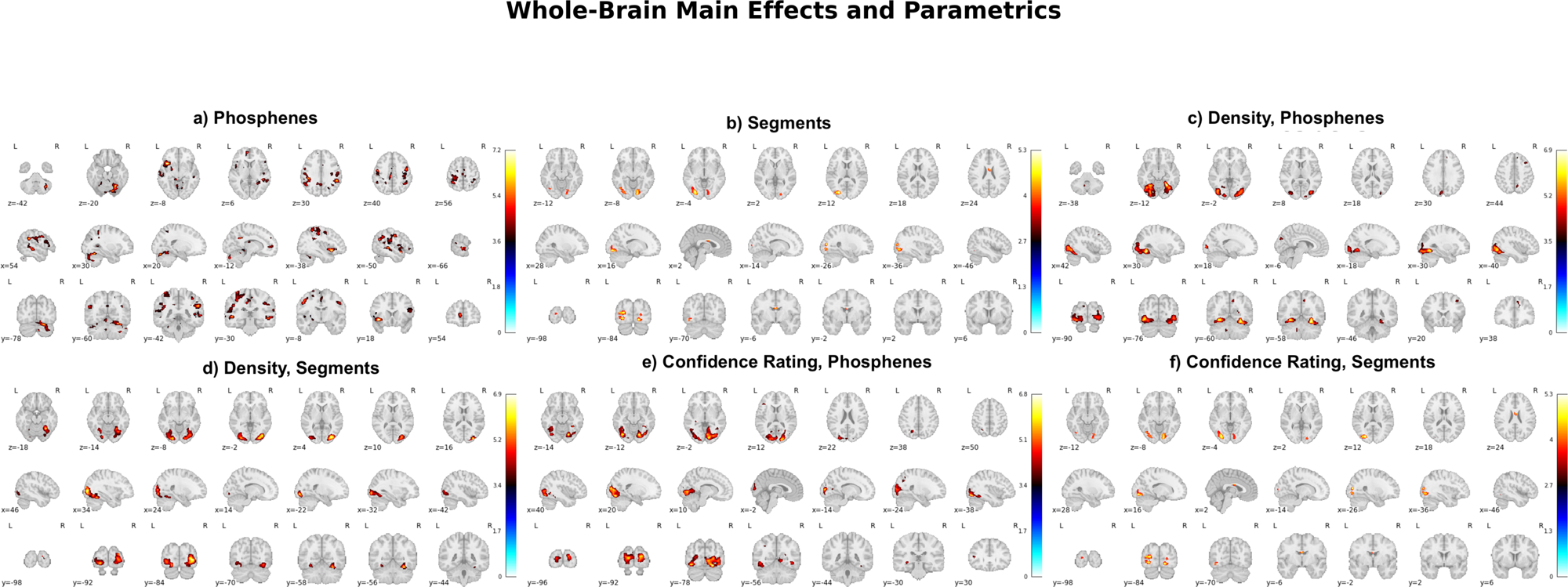
Whole brain results for different fragment conditions. a) Phosphene trials at fragmented stimulus presentation trigger responses in a large clusters centered on the middle and anterior cingulate, extending to the thalamus and pallidum. b) Curve segment trials shows a similar pattern to phosphene trials, albeit with more conservative spatial extents in the same significant regions. d) and c) Contrast maps for t-tests on fragment densities in both conditions yield significant responses in the occipital cortex. e) and f) Confidence in both fragment types shows a highly similar response pattern to density levels, showing the direct relationship between the two, as seen in behavioral measures, although confidence ratings for segment trials did not survive correction for multiple comparisons. Maps shown are thresholded at p = 0.05 FWE, cluster corrected (k = 25).

**Figure 3.**
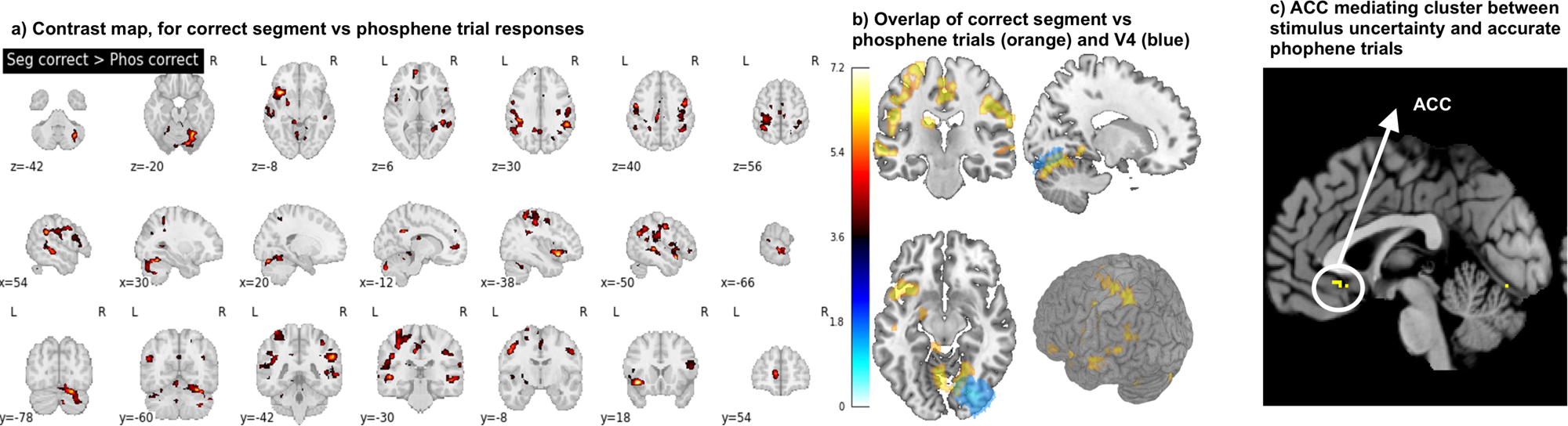
Whole brain results by fragment type at trial response. a) Correct curve segment trials tested against correct phosphene trials yields extensive responses in the occipital cortex encompassing area V4, as well bilateral middle temporal gyri, part of the visual ventral stream. In addition, this contrast yielded responses in the left insula, right inferior frontal gyrus and right parietal areas as well as a significant cluster in the anterior cingulate cortex. b) Statistical map of correct segments contrasted against correct phosphenes (orange) overlaid on the ROI of area V4, showing overlap (blue). c) Mediation analysis. Brain regions mediating the link between heightened uncertainty in phosphenes and correct performance include the anterior cingulate cortex (ACC).

**Table 1.** Contrast results at stimulus presentation onsets.

| Segments |  |  |  |  |  |  |  |
| --- | --- | --- | --- | --- | --- | --- | --- |
| p_FWE_corr_ | Cluster size k | p_FWE_corr__1 | T | x | y | z | Region_AAL3v1 |
| 0 | 1069 | 0.003 | 8.15 | -46 | -74 | 40 | Angular_L |
| 0 | 6602 | 0.004 | 7.95 | -6 | -8 | 2 | Thal_MDm_L |
| 0 | 1499 | 0.025 | 6.83 | 62 | -56 | 6 | Temporal_Mid_R |
| 0 | 723 | 0.131 | 5.86 | 0 | -28 | 42 | Cingulate_Mid_L |
| 0.005 | 460 | 0.316 | 5.3 | -54 | -22 | -14 | Temporal_Mid_L |
| 0.034 | 298 | 0.468 | 5.02 | -14 | -16 | -40 | Brainstem |
| Phosphenes |  |  |  |  |  |  |  |
| p_FWE_corr_ | Cluster size k | p_FWE_corr__1 | T | x | y | z | Region_AAL3v1 |
| 0 | 16085 | 0 | 10.32 | 6 | -2 | -4 | Accumbens, R, extending into ACC |
| 0 | 2012 | 0 | 10.04 | -42 | -68 | 50 | Angular_L, extending to MCC |
| 0 | 2617 | 0.001 | 8.9 | 48 | -42 | 2 | Temporal_Mid_R |
| 0 | 1716 | 0.034 | 6.8 | -52 | -22 | -20 | Temporal_Inf_L |
| 0.013 | 300 | 0.099 | 6.18 | -20 | -48 | 4 | Precuneus_L |
| 0.007 | 342 | 0.133 | 6.01 | 28 | -84 | -32 | Cerebellum_Crus1_R |
| 0.023 | 265 | 0.191 | 5.78 | -22 | -86 | -34 | Cerebellum_Crus2_L |
| Phosphenes, Density |  |  |  |  |  |  |  |
| p_FWE_corr_ | Cluster size k | p_FWE_corr__1 | T | x | y | z | Region_AAL3v1 |
| 0 | 2374 | 0.023 | 6.9 | 30 | -58 | -12 | Fusiform_R |
| 0 | 2534 | 0.048 | 6.47 | -30 | -60 | -10 | Fusiform_L |
| Segments, Density |  |  |  |  |  |  |  |
| p_FWE_corr_ | Cluster size k | p_FWE_corr__1 | T | x | y | z | Region_AAL3v1 |
| 0 | 2132 | 0.016 | 6.87 | 36 | -84 | 4 | Occipital_Mid_R |
| 0 | 1191 | 0.094 | 5.86 | -22 | -90 | -8 | Occipital_Inf_L |
| Phosphenes, Confidence |  |  |  |  |  |  |  |
| p_FWE_corr_ | Cluster size k | p_FWE_corr__1 | T | x | y | z | Region_AAL3v1 |
| 0 | 5390 | 0.03 | 6.79 | -38 | -82 | -10 | Occipital_Inf_L |

**Table 2.** Contrast results at response onsets.

| Segment Correct > Phosphene Correct |  |  |  |  |  |  |  |
| --- | --- | --- | --- | --- | --- | --- | --- |
| p_FWE | k | p_FWE | T | x | y | z | Region_AAL3v1 |
| 0 | 844 | 0.016 | 7.17 | -50 | 12 | -14 | Temporal_Pole_Sup_L |
| 0 | 1903 | 0.044 | 6.6 | 30 | -60 | -20 | Cerebellum_6_R |
| 0 | 2343 | 0.098 | 6.12 | 56 | -42 | 32 | SupraMarginal_R |
| 0 | 2731 | 0.126 | 5.97 | -44 | -34 | 32 | SupraMarginal_L |
| 0.003 | 436 | 0.14 | 5.91 | -50 | -30 | 0 | Temporal_Mid_L |
| 0.001 | 533 | 0.198 | 5.7 | 50 | -34 | -4 | Temporal_Mid_R |
| 0.001 | 536 | 0.372 | 5.28 | -6 | -26 | 40 | Cingulate_Mid_L |

#### ROI Analysis

To explicitly test our hypotheses regarding specific visual areas preferentially relating to specific fragment types, we examined those whole brain contrasts relating to different fragment trials that yielded significant responses using an ROI approach. Specifically, we tested contrast images on correct phosphene trials versus correct segment trials at response onset; phosphene trials alone, at stimulus onset; and segment trials alone, at stimulus onset. The *a priori* ROIs selected included: V1 (bilateral, ventral and dorsal); V4 (bilateral); and lateral occipital cortex (2 regions, bilateral). The first two regions are of direct interest to our hypothesis and the LOC was added as a region known to be implicated in object recognition. The specific ROIs used were taken from a probabilitic retinotopic atlas (L. Wang et al., 2015). Of the contrasts tested, we found that phosphene trials at stimulus presentation elicited significant responses in the right lateral occipital cortex (1 and 2) and bilaterally in V1 (ventral). Segment trials yielded no significant responses in any of the ROIs tested. Correct segment trials greater than correct phosphene trials however yielded significant responses in bilateral V1 (ventral) and V4 (Table 3).

**Table 3.** Mediation analysis on regions implicated in the relationship between stimulus uncertainty and accuracy.

| Peak | peak |  |  |  |
| --- | --- | --- | --- | --- |
| p(FWE-corr) | T | x | y | z |
| Phosphenes, stimulus presentation onset |  |  |  |  |
| roi14_R, lateral occipital cortex 2 |  |  |  |  |
| 0.034 | 4.53 | 50 | -74 | 28 |
| roi15_R, lateral occipital cortex 1 |  |  |  |  |
| 0.048 | 4.43 | 54 | -70 | 20 |
| roi1_R, V1 ventral |  |  |  |  |
| 0.020 | 4.91 | 48 | -44 | -14 |
| Phosphene Correct < Segment Correct, response onset |  |  |  |  |
| roi1_L, V1 ventral |  |  |  |  |
| 0.023 | 4.9 | -6 | -72 | -10 |
| roi1_R, V1 ventral |  |  |  |  |
| 0.009 | 5.36 | 12 | -76 | -12 |
| roi7_L, hV4 |  |  |  |  |
| 0.021 | 4.78 | -8 | -80 | -16 |
| roi7_R, hV4 |  |  |  |  |
| 0.002 | 5.94 | 14 | -68 | -10 |

#### Mediation Analysis

We next sought to identify brain areas involved in mediating the relationship between stimulus uncertainty, computed above, and correct object recognition for each fragment type. We found that correct responses in phosphene fragments under higher uncertainty were dependent on the anterior cingulate cortex (Figure 3c) and areas V1 and V4, while in segments, relied only on V4.

**Table 4.** Description of regression models constructed to examine the effects of decision-making variables and targets on behavioral responses.

| Model | Outcome Variable | Predictor Variables | Random Effects |
| --- | --- | --- | --- |
| Model 1<br>(binomial) | Accuracy | Fragment Type, Density, Uncertainty*, Fragment Type * Density, Fragment Type * Uncertainty, Reaction Time. | Participant, object ID |
| Model 2 | Confidence | Fragment Type, Density, Uncertainty*, Fragment Type * Density, Accuracy, Reaction Time | Participant, object ID |

**Table 5.** Results of mediation effect between uncertainty in visual stimulus and accuracy.

| Phosphenes, mediation *ab* effect |  |  |  |  |  |
| --- | --- | --- | --- | --- | --- |
| Name | x | y | z | voxels | maxstat |
| V1 | 0 | -78 | -8 | 4 | 7.1 |
| ACC | 2 | 26 | -6 | 26 | 9.97 |
| V1 | 2 | -100 | 4 | 1 | 5.91 |
| V1 | -8 | -104 | 8 | 2 | 8.96 |
| V1 | -2 | -102 | 8 | 1 | 20.76 |
| V4 | 32 | -94 | 12 | 6 | 8.01 |
| Segments, mediation *ab* effect |  |  |  |  |  |
| Name | x | y | z | voxels | maxstat |
| V4 | 14 | -68 | -6 | 3 | 7.19 |

##### Multivariate Pattern Analyis

Single-trial beta estimates were separated by fragment type to identify which predefined ROIs significantly predicted successful object recognition (correct vs. incorrect classification).In phosphene trials, the following regions predicted successful object recognition: right ACC (t=1.939, p=0.0334, d=0.42); left V1 (t=1.860, p=0.0388, d=0.41); left and right lateral occipital cortex 2 (t=1.882, p=0.0373, d=0.41; t =2.068, p=0.0259, d=0.45) and left and right occipital cortex 1 (t =3.336, p =0.0016, d=0.7; t(20)=2.430, p=0.0123, d=0.53). In segment trials, only left V4 predicted successful object recognition: t=2.169, p=0.0206, d=0.45). Together, these results highlight that successful object reocgnition from phosphenes recognition recruits an expanded occipital–frontal network, whereas object recognition from segments depends more on V4.

## Discussion

In this study, we queried three hypotheses relating to object recognition from sparse visual information. We asked 1) whether curve segments provide a more efficient basis for object recognition than low-level phosphenes; 2) if recognition from these different fragment types would recruit distinct visual areas, with V4 supporting segments and V1 supporting phosphenes; and 3) that successful recognition under uncertainty would depend on downstream cortical circuitry. Our behavioral results confirmed the first hypothesis: segments required lower fragment densities than phosphenes and yielded higher accuracy and confidence as fragment density increased. In line with the second hypothesis, correct segment-based recognition, relative to phosphenes, was associated with BOLD responses in area V4 and a more distributed occipito–temporo–parietal network, whereas phosphenes predominantly engaged V1 and lateral occipital cortex at stimulus onset. Whole-brain contrasts showed that correct recognition of segments, relative to phosphenes, evoked a more widespread network response. To further test whether these regions carried information predictive of successful recognition, we used multivariate pattern analysis to decode object recognition for each fragment type. The decoding results confirmed this functional dissociation: V1, LOC, and ACC predicted correct recognition from phosphenes, whereas only V4 predicted recognition from segments. To address the third hypothesis, mediation analyses were conducted to query whether correct recognition under uncertainty was supported by the ACC and visual cortex: for phosphenes, ACC (together with V1/V4) mediated successful object recognition, whereas this relationship was mediated primarily through V4 for segments, indicating that structured mid-level input reduces the need for ACC-based resolution of uncertainty and suggesting the latter reduce perceptual uncertainty and may rely more on sensory input rather than a downstream, cognitive mediation.

### Implications for Visual Prosthetics & Cortical Hierarchies (H1)

Our behavioral results suggest that eliciting curve segments, rather than phosphenes, may require fewer cortical stimulations, offering an empirically grounded means by which to simplify both the design and implantation of visual prostheses for the blind. Segments enabled accurate object recognition at lower density thresholds than phosphenes, confirming our first hypothesis (H1) that mid-level visual features provide a more efficient code for object recognition. This is in line with previous work showing that adding contour information to phosphenes improved letter recognition in simulated prosthetic vision (Kiral-Kornek et al., 2014). At the neural level, we found shared V1 responses for both fragment types but a unique V4 response for segments at the time of correct recognition, relative to phosphenes. The MVPA results refine this interpretation by showing that V4 at stimulus presentation uniquely carries information predictive of object recognition from segments. This finding lays the groundwork for exploratory stimulation of V4 in patients, to test whether curve segment perception can be artificially induced. Direct stimulation of ventral visual cortex in humans (Parvizi et al., 2012; Rangarajan et al., 2014) shows that perceptual outcomes depend on both cortical site and attentional state, implying that any V4-based prosthetic interface will require task- or context-dependent calibration. Previous efforts to artificially stimulate phosphenes in extrastriate areas have been successful in V2 and V3, and less promising in V5 and LOC (Schaeffner & Welchman, 2017). Studies focusing on V4 stimulation in macaques have been inconsistent with one finding no enhancement of V1 phosphene detection thresholds (Dagnino et al., 2015), and another, increased stimulus detection (Kienitz et al., 2022). Our data, in conjunction with very recent layer fMRI mapping of curvature-preference domains within human V4 (Zamboni et al., 2025), suggest this region encodes precisely the type of mid-level contour information that could be exploited by cortical stimulation to yield more coherent percepts than point-based inputs. Critically, V4 emerges as a promising alternative or complementary target to V1 for prosthetic development, offering the potential for sparser yet more informative stimulation patterns.

Beyond the visual hierarchy, our results highlight a mediating role for the anterior cingulate cortex (ACC). Both fragment types engaged the ACC at stimulus presentation, but mediation analysis showed that phosphene-based recognition relied more heavily on ACC contributions, consistent with its broader role in resolving uncertainty (Levinson et al., 2021; O’Reilly et al., 2013; Y. Wang et al., 2017) including visual uncertainty (Levinson et al., 2021). This downstream involvement may help explain why artificially induced phosphene perception is so variable across individuals: differences in the efficacy of cingulate-driven resolution may amplify variability in conscious phosphene experience (Phylactou et al., 2023) and further echoes reports from human stimulation work showing task-dependent perceptual modulation in ventral cortex (Parvizi et al., 2012), supporting the view that prosthetic percepts will necessarily interact with cognitive state and attentional control. By contrast, when contour fragments are encoded as mid-level segments, successful recognition can be mediated primarily within visual cortex, reducing the dependence on ACC and suggesting a more “locally solvable” perceptual problem for the brain.

### Theoretical Context & Model Interpretations (H2 & H3)

These results dovetail with predictive coding accounts in which ACC provides corrective or prior-based input when sensory evidence is poor or sparse (Alexander & Brown, 2014; Gold & Stocker, 2017; Kéri et al., 2004; Rushworth & Behrens, 2008). Phosphenes, being more ambiguous, appear to demand stronger ACC contributions, while segments reduce uncertainty by engaging mid-level visual areas more directly. The decoding results align with this framework: recognition-related information engages ACC and early visual cortex for phosphenes, but recruits only V4 for segments, consistent with a shift from prior-driven to evidence-driven inference. This interpretation is further supported by human fMRI evidence that V4 contributes to global shape integration and curvature tuning (Lawrence et al., 2023), and that contour-integration dynamics over time engage higher ventral areas beyond V1 (Kuai et al., 2017). Together with curvature-selective domains revealed in V4 (Pasupathy et al., 2020; Zamboni et al., 2025), these results converge on a mid-level mechanism by which contour continuity reduces uncertainty and stabilizes recognition(Choi et al., 2018) and further aligns with findings showing that local feedback from the extrastriate cortex to V1 is an important mechanism enabling object recognition under challenging conditions such as with fragmented, unclear or occluded objects (S. Chen et al., 2021; Wyatte et al., 2014). In predictive-coding terms, phosphene-based inputs provide weak and noisy likelihoods, forcing higher-order regions such as ACC to impose priors to stabilize perception, whereas segment-based inputs deliver richer mid-level evidence in V4, allowing recognition to proceed with less reliance on top-down information.

A further dimension to this interpretation comes from the confidence data. Confidence judgments correlated with activity in visual cortex, suggesting that metacognitive monitoring in this task is at least partly contained within the sensory system itself (Heereman et al., 2015) though other studies have found evidence of metacognitive tracking elsewhere in the brain (Balsdon et al., 2021; Molenberghs et al., 2016). This has direct implications for prosthetics: artificial stimulation must not only reach an expected perceptual threshold but also be consciously registered in the individual. Confidence ratings offer a behavioral window onto perceptual quality, an index that could guide calibration in closed-loop prosthetic systems. If a percept is consistently accompanied by low confidence, even if objectively “seen” as confirmed by neuroimaging responses in the visual cortex, it may fail to provide functional utility (Meuwese et al., 2014; Michel et al., 2024). Future closed-loop prosthetic systems might incorporate confidence-like feedback or adaptive encoding to maintain perceptual stability across cognitive states. Our findings suggest that such calibration should prioritize stimulation patterns that naturally support higher confidence via robust mid-level encoding, rather than merely increasing overall stimulation intensity or electrode count.

### Limitations & Future Directions

Our study has several limitations. While fMRI provides insight into brain mechanisms, it still falls short of causal explanations. Future work should employ causal interventions such as TMS or intracortical stimulation to confirm whether segment-like percepts can be elicited by stimulating V4. Second, the generalizability of our findings to other fragment types remains to be determined; the perceptual efficiency of segments may depend on their specific structure. Finally, although V4 stimulation offers theoretical advantages, clinical translation faces challenges including surgical access, stimulation thresholds, and the need for precise electrode targeting. These challenges parallel those faced in existing intracortical prosthesis trials, where electrode stability and mapping drift remain unresolved issues (Oswalt et al., 2021). Addressing these challenges will require integrating causal stimulation studies with computational modeling approaches capable of predicting perceptual outcomes from cortical dynamics and explicitly incorporating mid-level representational constraints derived from human neuroimaging and behavior.

### Broader Relevance

These results inform the next generation of visual prostheses. Existing phosphene-based devices rely on dense V1 stimulation, yet often yield variable and unstable percepts. By showing that curve segments engage V4 and reduce the need for ACC-driven resolution, our work suggests a new avenue for prosthetic design: stimulation strategies that approximate mid-level features rather than isolated phosphenes. Moreover, our findings can enrich computational prosthetic simulators by incorporating system-level constraints such as ACC stabilization. Recent computational pipelines now enable biologically plausible, differentiable simulators that optimize stimulation parameters end-to-end (Fine & Boynton, 2024; van der Grinten et al., 2024). Incorporating behavioral priors favoring contour continuity and curvature structure into these simulators could bias optimization toward activation patterns that exploit cortical organization for perceptual coherence. Similarly, human-in-the-loop deep learning frameworks for prosthetic encoding (Granley et al., 2023) could integrate these mid-level priors to personalize stimulation patterns that minimize crowding and energy requirements. Thus the “segment advantage” identified here not only clarifies the processes supporting recognition under sparse input but also provides a computational constraint for the emerging class of virtual-patient simulators and adaptive encoders aimed at restoring functional vision pointing toward cortical prostheses that elicit curves rather than dots to achieve structured, stable artificial sight.

## Methods

### Population

The study was conducted at the Laboratoire de Recherche en Neuroimagerie (Centre Hospitalier Universitaire Vaudois, Lausanne, Switzerland) and approved by the Commission d’éthique de la Recherche, Vaud. Participants were recruited from the general population and provided with study information two days prior to the session. On the day of testing, they reviewed and signed informed consent. Inclusion criteria were age 18–45 and good general health; exclusion criteria were self-reported psychiatric or neurological diagnoses, current or recent psychotropic medication, claustrophobia, and metal or other implants incompatible with MRI. Visual acuity was screened with the Fribourg Visual Acuity test (Landolt C); all participants met the minimum requirement of logMAR ≤ 0.3 (20/40 Snellen equivalent). After screening, the experimenter explained MRI procedures and task instructions before participants entered the scanner. Data were collected between April and May 2024.

### Cohort

The study recruited 27 participants, of which 24 were retained. Three participants were excluded due to problems with the MR headcoil during acquisition. The cohort included 24 participants (11females) that completed the experiment, with 23 included in image analysis. One participant had to be excluded from imaging analysis for faulty timing recordings. As a control for performance, we verified that participants made no more than 1 error of object recognition for contour and RGB trials. All participants passed this threshold.

### MRI acquisition

MRI data were acquired on a 3T Siemens Prisma scanner (Siemens, Erlangen, Germany). For each participant, functional data was acquired using a custom-made dual-echo 2D gradient-echo EPI sequence with echo times 17.4 and 35.2ms. The number of slices was 40 and the repetition time was TR = 2.48 s. The radio-frequency excitation flip angle was 90°, in-plane resolution was 3mm. Two functional scans of approximately 20 min each were acquired during task performance. B_0_-field mapping data were acquired using a 2D gradient-recalled sequence (TR = 700 ms, TE = 7.38 ms, flip angle = 80°, 64 × 64 × 64 voxels, ≈ 3 mm isotropic) to correct for geometric distortions in the EPI images. T1-weighted anatomical data scan was also acquired using a 3D MPRAGE sequence with the following parameters: TR = 2000 ms, TE = 2.39 ms, flip angle = 9°; matrix = 232 × 256 × 176; voxel size ≈ 1 mm isotropic.

### Stimulus generation procedure

Experimental stimuli consisted in fragmented images containing phosphenes or curved segments along object’s contours. We followed the fragmentation procedure used in (Lonnqvist et al., 2025) and (Scialom et al., 2024) . The fragmentation algorithm takes an RGB image as input, extracts its contours and places phosphenes and curved segments at the same pixel coordinates along the object’s starting outlines. To make both stimulus types comparable, phosphene widths were equal to segment lengths, spanning approximately 0.27 arcdeg. A density of 100% fragments corresponds to the maximum number of fragments that can outline the object’s contours. The set of segments was taken from (Mineault et al., 2013)), as it triggered specific V4 activity in macaque monkeys. Details of the algorithm are described in (Scialom et al., 2024).

### Task procedure

Prior to behavioral data collection and following the acquisition of the T1-MPRAGE anatomical scan, participants performed 5 practice trials of the task in the scanner. These trials employed stimuli that were not used in the experimental task. Participants were free to communicate with the experimenter via the microphone during the practice session in case of questions. Physiological measures (cardiac and respiratory) were recorded with a BioPac system. Once participants confirmed their readiness to perform the task, the first of two-trial series was launched.

In total, participants performed a total of 144 trials divided into two blocks. Stimuli were taken from the Bank of Standardized Stimuli and consisted of common objects (Brodeur et al., 2010). Eight distinct objects were selected as target stimuli and an additional 12 were selected as decoys. The 8 target stimuli were fragmented into either dot (phosphene condition) or curve segment (segment condition) outlines at the following fragment density levels: 12%, 16%, 21%, 27%, 35%, 46%, 59% and 77%, following an exponential fit previously used in (Torfs et al., 2010).

In each block, 4 distinct target objects were presented, in a randomized fashion across objects and fragment condition, at increasing fragment density levels (8), followed by the same 4 objects shown in full (white) contour on a black background; and then the same 4 target objects shown in full RGB. Each trial began with the presentation of a black fixation cross on a grey background. Participants were asked to perform an object recognition task. At each trial, an image of an object (15 x 15 visual degrees; cue) was first briefly presented (120 ms), followed by a black fixation cross on a grey background (6 s). Then, participants were shown a 3 * 3 grid of different objects in full color at a size of 5 * 5 degrees. This grid contained 5 decoys and 4 possible target stimuli, including one image of object category that was previously shown (Target). While the target was the same object as the cue, it was not the same image, but an example from a given image category (e.g. “boots”). For instance, the cue may have been a pair of glasses oriented towards the bottom left of the screen, and the target, oriented towards the bottom right. Participants were asked to select the previously shown object from the grid using a button response box. Following their selection, they were shown another black fixation cross on a grey background for approximately 6 s. Finally, participants were asked to provide a metacognitive confidence report on a 1-10 scale (Figure 4a). All images were shown with white fragments on a black background, except for the RGB images, which were shown on a grey background. All code was written in Python using the PsychoPy presentation library.

**Figure 4:**
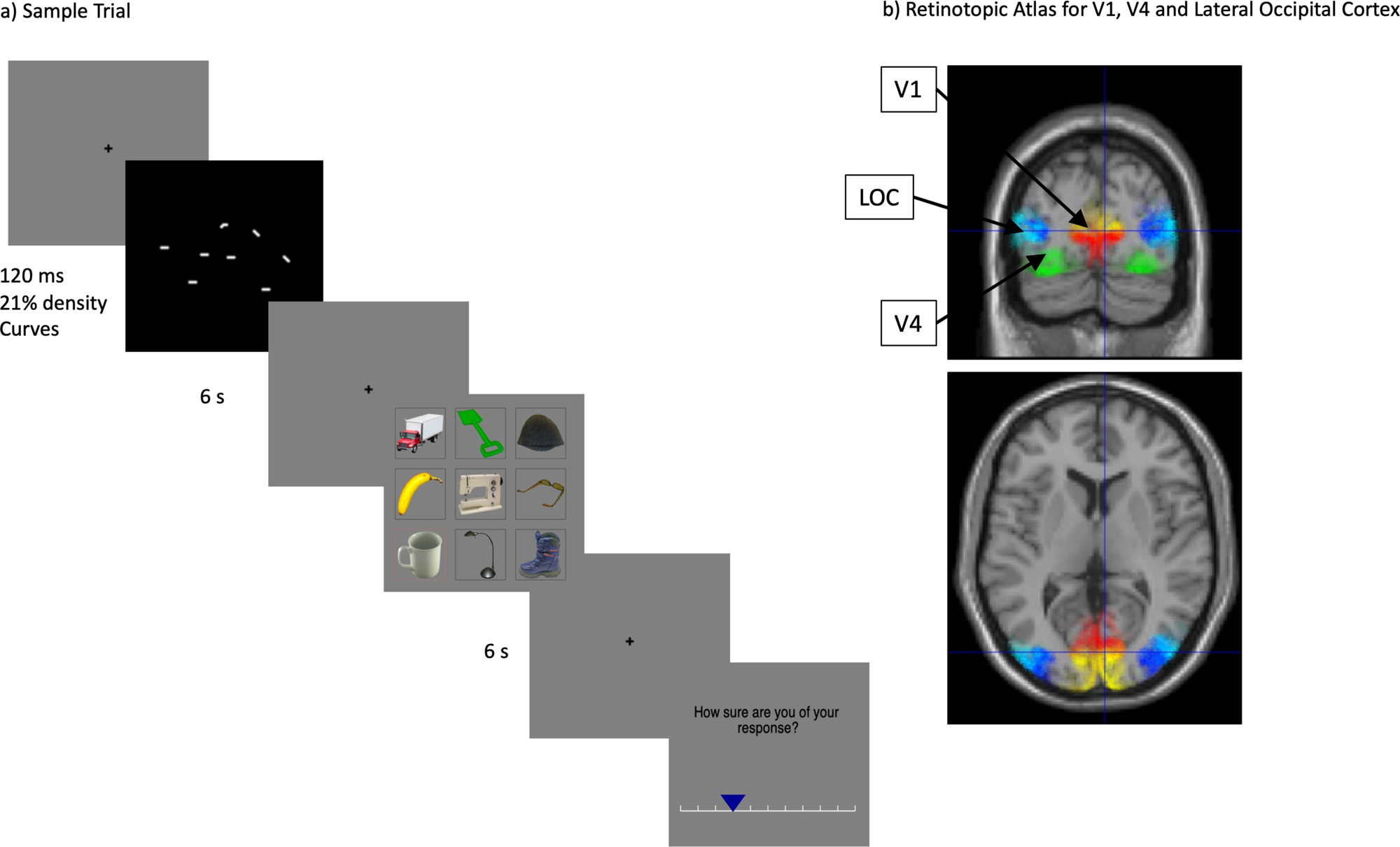
a) Sample trial. Participants were briefly shown (120 ms) an image, composed of either white curve segments or phosphenes (dots) on a black background. The image here shows a pair of glasses at a fragment density of 21%. Participants are then shown a 3*3 grid of objects, including target stimuli as well as decoy objects and asked to navigate the grid to find the matching object. Matching objects always varied in their identity. Participants were then asked to rate their confidence in their choice. b) Retinotopic map of regions selected for ROI analysis, inluding lateral occipital cortex (LOC) and areas V1 and V4 ((L. Wang et al., 2015).

## Data Analysis

### Behavioral

To characterize perceptual sensitivity, individual psychometric curves were fit separately for phosphenes and segments. Percent correct was computed at each fragment density (8 levels), and individual subject data were fit with a logistic function, yielding slope (*k*) and point of subjective equality (PSE) parameters.

Trial-wise perceptual uncertainty was estimated by computing the entropy of the predicted probability of a correct response (*p_c_*), derived from each subject’s logistic fit. This produced an uncertainty measure that varied with stimulus density. These trial-level values were used in subsequent analyses.

Mixed-effects regression models were then fit to test how 1) accuracy and 2) confidence were predicted by density, uncertainty, and stimulus type. Models included random intercepts for subject (and object identity when available), and binomial link functions for accuracy (Table 1).

### Neuroimaging analysis

Neuroimaging analyis was performed using a mass univariate analysis in SPM12 (Wellcome Centre for Human Neuroimaging, UCL, UK www.fil.ion.ucl.ac.uk/spm). The generalized linear model was designed based on the behavioral models above, adapted for fMRI. The preprocessing pipeline applied to the functional MR datasets is described below.

### Preprocessing

Functional MR volumes were first converted to niftii format from dicom. The two echo images for each volume of the fMRI time-series were combined using simple summation (Poser et al., 2006). Images were then slice-time corrected to the middle slice. Voxel displacement maps were computed from the B_0_-field mapping data using SPM’s Fieldmap toolbox (Andersson et al., 2001) and inserted into the realign and unwarp step of SPM (6; 3 translational and 3 rotational). Individual T1w anatomical volumes were then co-registered to the mean functional images, followed by tissue classification and spatial registration to standard Montreal Neurological Institute (MNI) space using SPM12s “unified segmentation”. The resulting parameters (forward deformations) were then applied to functional volumes, before applying a Gaussian 3D smoothing with 8mm full-width-at-half-maximum smoothing kernel. The resulting preprocessed images were used for statistical analysis.

### Voxelwise analysis

We estimated subject-level event-related GLMs in SPM12. Each participant contributed two EPI runs (TR = 2.48 s) modeled separately within a single design matrix. Regressors were convolved with the canonical hemodynamic response function. Low-frequency drifts were attenuated with a 128 s high-pass filter and serial correlations were modeled using FAST. Analyses were restricted to voxels within SPM’s intracranial volume mask; the implicit baseline served as the reference.

The first-level GLM included the following event onsets: (1) cue onsets corresponding to cue image presentation; (2) button-press responses; and (3) confidence reports. Each event type was modeled as a zero-duration (stick) function. Cue onsets were modeled separately for *phosphene* and *segment* fragment types, each parametrically modulated by (i) the fragment density value and (ii) its associated uncertainty, defined as the entropy value computed from behavioral data. RGB and contour trials were modeled as control conditions without modulators. Response events were modeled separately for phosphene and segment trials and were parametrically modulated by accuracy and reaction time. Confidence events were likewise modeled separately for the two fragment types and parametrically modulated by the confidence rating and reaction time. Within each condition, serial orthogonalization of parametric modulators was retained, such that variance was first attributed to the earlier modulator before the subsequent one.

Nuisance predictors included the six rigid-body head-motion parameters and 18 physiological regressors derived using RETROICOR (via the TAPAS PhysIO toolbox; Frässle et al., 2021) to capture respiration, cardiac phase, and interaction terms.

Model estimation used SPM’s “Classical” ordinary least squares. The mass univariate analyses tested effects associated with (i) phosphene trial onsets, (ii) segment trial onsets, and (iii) their direct contrasts, as well as the corresponding parametric effects of density, accuracy, and confidence. Additional contrasts pooled across fragment types to examine main effects of correct responses, density, and confidence, and compared fragmented-image trials with contour and RGB control trials. Statistical inference at the group level used a cluster-forming threshold of voxel-wise *p* < 0.001 (uncorrected, *k* = 25) and a family-wise error (FWE) correction at *p* < 0.05, following current best-practice recommendations (*Eklund et al.*, 2016).

### ROI analysis

Because the research question focuses on visual areas involved in object recognition, four *a priori* regions of interest (ROIs) were selected for region.based analysis: V1, V4, and the lateral occipital complex (LOC) in the visual cortex (L. Wang et al., 2015), in addition to the anterior cingulate cortex. Within each ROI, results were considered significant at *p* < 0.05, voxel-level FWE corrected using small-volume correction (Figure 4b).

### Mediation Analysis

To determine which brain regions mediate the relationship between the empirically derived uncertainty of a stimulus at a given fragment type and density level and correct object recognition. Mediation was conducted in Matlab using the CanLab mediation toolbox (with a bootstrapping procedure (5000 samples) within a mask confined to regions of interest in the visual cortex (V1, V4 and LOC) and the anterior cingulate cortex,

Single-trial cue responses for mediation were estimated with a beta-series (LS-A) GLM in SPM12. For each run, we modeled every cue onset as a separate stick regressor (64 trials/run; 0-s duration), together with shared regressors for response onsets (parametric modulators: accuracy, RT), confidence reports (parametric modulators: rating, RT), and control cue types (RGB, contours). Nuisance regressors comprised 6 rigid-body motion parameters and 18 physiological terms (RETROICOR), and a 128-s high-pass filter was applied; serial correlations were modeled with FAST. This yielded 128 cue betas per subject (64 per run).

Mediation analysis used Canlab’s multilevel brain mediation toolbox on *a priori* ROIs (mask comprising ACC, V1, V4, and LOC). The *X* variable was trial-wise perceptual uncertainty (Shannon entropy from each participant’s psychometric function); the *Y* variable was trial-wise accuracy (0/1); the mediator *M* was the vector of single-trial cue betas for each subject within the mask. Models were fit with random subject intercepts and bias-corrected bootstrap for inference (5000 samples); family-wise error within the mask was controlled by FDR and thresholding at q <0.05 on the voxelwise *p*-maps for the indirect path (*ab*). Fragment-type–specific models were obtained by gating *X* to phosphene or segment trials, with the same *M* and *Y*. Density was included as a covariate in all models.

### Multivariate Pattern Analysis (MVPA)

Multivariate pattern analysis (MVPA) tests were performed in addition to the univariate analysis to examine whether distributed voxel activation patterns encode behaviorally relevant information. MVPA thus provides more fine-grained information relative to the GLM, which focused on average signal changes in the brain. MVPA therefore allows us to identify whether a pattern of responses predicts behavior even when standard univariate analyses show no mean activation differences.

In this study, MVPA was used to test whether trial-by-trial activity in the above set of a priori ROIs predicted successful object recognition, separately for phosphene and segment trials. For each participant, we started from the single-trial GLM used for the mediation analysis above, where the beta images were extracted for single-trial cue-onset regressors. For every trial we extracted fragment type (phosphenes versus segments) and a binary recognition accuracy label (correct vs incorrect). Trials from blocks 1 and 2 were used as separate cross-validation chunks. ROIs were taken from the probabilistic atlas used in the univariate analyses (including bilateral ACC, V1, V4 and LOCs 1 and 2).

Decoding was performed with The Decoding Toolbox (TDT v3.994; SPM12 backend). Within each fragment type, the classifier was trained to discriminate correct from incorrect trials using a leave-one-session-out cross-validation scheme, and we extracted balanced accuracy minus chance as the scalar decoding metric from each ROI. For each condition and ROI we then tested, across subjects, whether accuracy-minus-chance was significantly greater than zero using one-sample, one-tailed t-tests. This yielded, for each ROI and fragment type, group-level estimates of above-chance that we use to identify which regions reliably predict correct object recognition in phosphenes and in segments.

## Acknowledgments

We thank Jenny Schroeder for her instrumental contribution in neuroimaging data acquisition. We thank Prof. Udo Ernst and Dr. David Rotermund for providing the experimental stimuli used in the study. LLK, ES, MH and BD were funded by ERA_NET NEURON JTC2020: iSEE (https://www.neuron-eranet.eu/projects/I-See/). LLK was also funded by the Faculté de Biologie et de Médecine, Université de Lausanne, Transition grant. AL is funded by the Swiss National Science Foundation (https://data.snf.ch/grants) project grants no. Grant Nos. 320030_184784 and CR00I5-235940. BD is also funded by the Swiss National Science Foundation (https://data.snf.ch/grants) project grants no. 213595, 32003B_135679, 32003B_159780, 324730_192755 and CRSK-3_190185) and JTC2023-ELSA: BrainTree project (https://www.neuron-eranet.eu/projects/BRAINTREE/) and the InnoSuisse Flagship Swiss brAInHealth project (https://www.innosuisse.admin.ch/en/ongoing-flagships). The Laboratory for Research in Neuroimaging—LREN is very grateful to the Roger De Spoelberch (https://www.fondation-roger-de-spoelberch.ch/) for their generous financial support.

## CRediT

LLK: Conceptualization ; Project Administration; Investigation; Methodology; Formal Analysis; Writing – Original Draft Preparation;

ES: Conceptualization ; Writing – Review & Editing

AL: Methodology; Investigation; Writing – Review & Editing

MH: Writing – Review & Editing; Supervision; Funding Acquisition

BD: Writing – Review & Editing; Supervision; Funding Acquisition

## Declaration of generative AI and AI-assisted technologies in the manuscript preparation process

During the preparation of this work the author(s) used ChatGPT in order to enhance the prose and structural flow of the manuscript. After using this tool, the author(s) reviewed and edited the content as needed and take full responsibility for the content of the published article.

## References

Alexander, W. H., & Brown, J. W. (2014). A general role for medial prefrontal cortex in event prediction. 8(July), 1–11. 10.3389/fncom.2014.00069

Andersson, J. L. R., Hutton, C., Ashburner, J., Turner, R., & Friston, K. (2001). Modeling Geometric Deformations in EPI Time Series. NeuroImage, 13(5), 903–919. 10.1006/nimg.2001.0746

Balsdon, T., Mamassian, P., & Wyart, V. (2021). Separable neural signatures of confidence during perceptual decisions. eLife, 10, e68491. 10.7554/eLife.68491

Biederman, I. (1987). Recognition-by-components: A theory of human image understanding. Psychological Review, 94(2), 115–147. 10.1037/0033-295X.94.2.115

Borda, E., & Ghezzi, D. (2022). Advances in visual prostheses: Engineering and biological challenges. Progress in Biomedical Engineering, 4(3), 032003. 10.1088/2516-1091/ac812c

Bosking, W. H., Oswalt, D. N., Foster, B. L., Sun, P., Beauchamp, M. S., & Yoshor, D. (2022). Percepts evoked by multi-electrode stimulation of human visual cortex. Brain Stimulation, 15(5), 1163–1177. 10.1016/j.brs.2022.08.007

Brodeur, M. B., Dionne-Dostie, E., Montreuil, T., & Lepage, M. (2010). The Bank of Standardized Stimuli (BOSS), a New Set of 480 Normative Photos of Objects to Be Used as Visual Stimuli in Cognitive Research. PLOS ONE, 5(5), e10773. 10.1371/journal.pone.0010773

Carlson, E. T., Rasquinha, R. J., Zhang, K., & Connor, C. E. (2011). A Sparse Object Coding Scheme in Area V4. Current Biology, 21(4), 288–293. 10.1016/j.cub.2011.01.013

Chen, S. C., Suaning, G. J., Morley, J. W., & Lovell, N. H. (2009). Simulating prosthetic vision: I. Visual models of phosphenes. Vision Research, 49(12), 1493–1506. 10.1016/j.visres.2009.02.003

Chen, S., Weidner, R., Zeng, H., Fink, G. R., Müller, H. J., & Conci, M. (2021). Feedback from lateral occipital cortex to V1/V2 triggers object completion: Evidence from functional magnetic resonance imaging and dynamic causal modeling. Human Brain Mapping, 42(17), 5581–5594. 10.1002/hbm.25637

Choi, H., Pasupathy, A., & Shea-Brown, E. (2018). Predictive Coding in Area V4: Dynamic Shape Discrimination under Partial Occlusion. Neural Computation, 30(5), 1209–1257. 10.1162/NECO_a_01072

Dagnino, B., Gariel-Mathis, M.-A., & Roelfsema, P. R. (2015). Microstimulation of area V4 has little effect on spatial attention and on perception of phosphenes evoked in area V1. Journal of Neurophysiology, 113(3), 730–739. 10.1152/jn.00645.2014

Erickson-Davis, C., & Korzybska, H. (2021). What do blind people “see” with retinal prostheses? Observations and qualitative reports of epiretinal implant users. PLOS ONE, 16(2), e0229189. 10.1371/journal.pone.0229189

Fernández, E., Alfaro, A., Soto-Sánchez, C., Gonzalez-Lopez, P., Lozano, A. M., Peña, S., Grima, M. D., Rodil, A., Gómez, B., Chen, X., Roelfsema, P. R., Rolston, J. D., Davis, T. S., & Normann, R. A. (2021). Visual percepts evoked with an intracortical 96-channel microelectrode array inserted in human occipital cortex. The Journal of Clinical Investigation, 131(23), e151331. 10.1172/JCI151331

Fine, I., & Boynton, G. M. (2024). A virtual patient simulation modeling the neural and perceptual effects of human visual cortical stimulation, from pulse trains to percepts. Scientific Reports, 14(1), 17400. 10.1038/s41598-024-65337-1

Frässle, S., Aponte, E. A., Bollmann, S., Brodersen, K. H., Do, C. T., Harrison, O. K., Harrison, S. J., Heinzle, J., Iglesias, S., Kasper, L., Lomakina, E. I., Mathys, C., Müller-Schrader, M., Pereira, I., Petzschner, F. H., Raman, S., Schöbi, D., Toussaint, B., Weber, L. A., … Stephan, K. E. (2021). TAPAS: An Open-Source Software Package for Translational Neuromodeling and Computational Psychiatry. Frontiers in Psychiatry, 12. 10.3389/fpsyt.2021.680811

Gold, J. I., & Stocker, A. A. (2017). Visual Decision-Making in an Uncertain and Dynamic World. Annual Review of Vision Science, 3(Volume 3, 2017), 227–250. 10.1146/annurev-vision-111815-114511

Granley, J., Fauvel, T., Chalk, M., & Beyeler, M. (2023). Human-in-the-Loop Optimization for Deep Stimulus Encoding in Visual Prostheses. Advances in Neural Information Processing Systems, 36, 79376–79398.

Kéri, S., Decety, J., Roland, P. E., & Gulyás, B. (2004). Feature uncertainty activates anterior cingulate cortex. Human Brain Mapping, 21(1), 26–33. 10.1002/hbm.10150

Kienitz, R., Kouroupaki, K., & Schmid, M. C. (2022). Microstimulation of visual area V4 improves visual stimulus detection. Cell Reports, 40(12), 111392. 10.1016/j.celrep.2022.111392

Kiral-Kornek, F. I., O Sullivan-Greene, E., Savage, C. O., McCarthy, C., Grayden, D. B., & Burkitt, A. N. (2014). Improved visual performance in letter perception through edge orientation encoding in a retinal prosthesis simulation. Journal of Neural Engineering, 11(6), 066002. 10.1088/1741-2560/11/6/066002

Kuai, S.-G., Li, W., Yu, C., & Kourtzi, Z. (2017). Contour Integration over Time: Psychophysical and fMRI Evidence. *Cerebral Cortex (New York*, N.Y.: 1991), *27*(5), 3042–3051. 10.1093/cercor/bhw147

Lawrence, S. J. D., Zamboni, E., Vernon, R. J. W., Gouws, A. D., Wade, A. R., & Morland, A. B. (2023). The Emergence of Tuning to Global Shape Properties of Radial Frequency Patterns in the Ventral Visual Pathway. The Journal of Neuroscience: The Official Journal of the Society for Neuroscience, 43(29), 5378–5390. 10.1523/JNEUROSCI.2237-22.2023

Levinson, M., Podvalny, E., Baete, S. H., & He, B. J. (2021). Cortical and subcortical signatures of conscious object recognition. Nature Communications, 12(1), 2930. 10.1038/s41467-021-23266-x

Liu, X., Chen, P., Ding, X., Liu, A., Li, P., Sun, C., & Guan, H. (2022). A narrative review of cortical visual prosthesis systems: The latest progress and significance of nanotechnology for the future. Annals of Translational Medicine, 10(12), 716. 10.21037/atm-22-2858

Lonnqvist, B., Scialom, E., Gokce, A., Merchant, Z., Herzog, M. H., & Schrimpf, M. (2025). *Contour Integration Underlies Human-Like Vision* (arXiv:2504.05253). arXiv. 10.48550/arXiv.2504.05253

Meuwese, J. D. I., van Loon, A. M., Lamme, V. A. F., & Fahrenfort, J. J. (2014). The subjective experience of object recognition: Comparing metacognition for object detection and object categorization. *Attention, Perception*, & Psychophysics, 76(4), 1057–1068. 10.3758/s13414-014-0643-1

Michel, M., Gao, Y., Mazor, M., Kletenik, I., & Rahnev, D. (2024). When visual metacognition fails: Widespread anosognosia for visual deficits. Trends in Cognitive Sciences, 28(12), 1066–1077. 10.1016/j.tics.2024.09.003

Mineault, P. J., Zanos, T. P., & Pack, C. C. (2013). Local field potentials reflect multiple spatial scales in V4. Frontiers in Computational Neuroscience, 7. 10.3389/fncom.2013.00021

Molenberghs, P., Trautwein, F.-M., Böckler, A., Singer, T., & Kanske, P. (2016). Neural correlates of metacognitive ability and of feeling confident: A large-scale fMRI study. Social Cognitive and Affective Neuroscience, 11(12), 1942–1951. 10.1093/scan/nsw093

O’Reilly, J. X., Schuffelgen, U., Cuell, S. F., Behrens, T. E. J., Mars, R. B., & Rushworth, M. F. S. (2013). Dissociable effects of surprise and model update in parietal and anterior cingulate cortex. Proceedings of the National Academy of Sciences, 110(38), E3660–E3669. 10.1073/pnas.1305373110

Oswalt, D., Bosking, W., Sun, P., Sheth, S. A., Niketeghad, S., Salas, M. A., Patel, U., Greenberg, R., Dorn, J., Pouratian, N., Beauchamp, M., & Yoshor, D. (2021). Multi-electrode stimulation evokes consistent spatial patterns of phosphenes and improves phosphene mapping in blind subjects. Brain Stimulation, 14(5), 1356–1372. 10.1016/j.brs.2021.08.024

Parvizi, J., Jacques, C., Foster, B. L., Witthoft, N., Rangarajan, V., Weiner, K. S., & Grill-Spector, K. (2012). Electrical stimulation of human fusiform face-selective regions distorts face perception. The Journal of Neuroscience: The Official Journal of the Society for Neuroscience, 32(43), 14915–14920. 10.1523/JNEUROSCI.2609-12.2012

Pasupathy, A., Popovkina, D. V., & Kim, T. (2020). Visual Functions of Primate Area V4. Annual Review of Vision Science, 6, 363–385. 10.1146/annurev-vision-030320-041306

Phylactou, P., Traikapi, A., & Konstantinou, N. (2023). One in four people fail to perceive phosphenes during early visual cortex transcranial magnetic stimulation. *Brain Stimulation: Basic*, Translational, and Clinical Research in Neuromodulation, 16(1), 23–24. 10.1016/j.brs.2022.12.012

Poser, B. A., Versluis, M. J., Hoogduin, J. M., & Norris, D. G. (2006). BOLD contrast sensitivity enhancement and artifact reduction with multiecho EPI: Parallel-acquired inhomogeneity-desensitized fMRI. Magnetic Resonance in Medicine, 55(6), 1227–1235. 10.1002/mrm.20900

Rangarajan, V., Hermes, D., Foster, B. L., Weiner, K. S., Jacques, C., Grill-Spector, K., & Parvizi, J. (2014). Electrical stimulation of the left and right human fusiform gyrus causes different effects in conscious face perception. The Journal of Neuroscience: The Official Journal of the Society for Neuroscience, 34(38), 12828–12836. 10.1523/JNEUROSCI.0527-14.2014

Roe, A. W., Chelazzi, L., Connor, C. E., Conway, B. R., Fujita, I., Gallant, J. L., Lu, H., & Vanduffel, W. (2012). Toward a Unified Theory of Visual Area V4. Neuron, 74(1), 12–29. 10.1016/j.neuron.2012.03.011

Rushworth, M. F. S., & Behrens, T. E. J. (2008). Choice, uncertainty and value in prefrontal and cingulate cortex. Nature Neuroscience, 11(4), 389–397. 10.1038/nn2066

Sanchez-Garcia, M., Martinez-Cantin, R., & Guerrero, J. J. (2018). Structural and object detection for phosphene images (arXiv:1809.09607). arXiv. 10.48550/arXiv.1809.09607

Schaeffner, L. F., & Welchman, A. E. (2017). Mapping the visual brain areas susceptible to phosphene induction through brain stimulation. Experimental Brain Research, 235(1), 205–217. 10.1007/s00221-016-4784-4

Scialom, E., Ernst, U. A., Rotermund, D., & Herzog, M. H. (2024). Minimizing the number of phosphenes required for object recognition under prosthetic vision (p. 2024.09.26.614440). bioRxiv. 10.1101/2024.09.26.614440

Shiozaki, H. M., Tanabe, S., Doi, T., & Fujita, I. (2012). Neural Activity in Cortical Area V4 Underlies Fine Disparity Discrimination. Journal of Neuroscience, 32(11), 3830–3841. 10.1523/JNEUROSCI.5083-11.2012

Torfs, K., Panis, S., & Wagemans, J. (2010). Identification of fragmented object outlines: A dynamic interplay between different component processes. Visual Cognition, 18(8), 1133–1164. 10.1080/13506281003693593

van der Grinten, M., de Ruyter van Steveninck, J., Lozano, A., Pijnacker, L., Rueckauer, B., Roelfsema, P., van Gerven, M., van Wezel, R., Güçlü, U., & Güçlütürk, Y. (2024). Towards biologically plausible phosphene simulation for the differentiable optimization of visual cortical prostheses. eLife, 13, e85812. 10.7554/eLife.85812

Wang, L., Mruczek, R. E. B., Arcaro, M. J., & Kastner, S. (2015). Probabilistic Maps of Visual Topography in Human Cortex. *Cerebral Cortex (New York*, N.Y.: 1991), *25*(10), 3911–3931. 10.1093/cercor/bhu277

Wang, Y., Ma, N., He, X., Li, N., Wei, Z., Yang, L., Zha, R., Han, L., Li, X., Zhang, D., Liu, Y., & Zhang, X. (2017). Neural substrates of updating the prediction through prediction error during decision making. NeuroImage, 157, 1–12. 10.1016/j.neuroimage.2017.05.041

Wei, H., Dong, Z., & Wang, L. (2018). V4 shape features for contour representation and object detection. Neural Networks, 97, 46–61. 10.1016/j.neunet.2017.09.010

Winawer, J., & Witthoft, N. (2015). Human V4 and ventral occipital retinotopic maps. Visual Neuroscience, 32, E020. 10.1017/S0952523815000176

Wyatte, D., Jilk, D. J., & O’Reilly, R. C. (2014). Early recurrent feedback facilitates visual object recognition under challenging conditions. Frontiers in Psychology, 5. 10.3389/fpsyg.2014.00674

Zamboni, E., Watson, I., Stirnberg, R., Huber, L., Formisano, E., Goebel, R., Kennerley, A. J., & Morland, A. B. (2025). Mapping curvature domains in human V4 using CBV-sensitive layer-fMRI at 3T. Frontiers in Neuroscience, 19. 10.3389/fnins.2025.1537026

